# Genome Wide Characterization, Identification And Expression Analysis Of *Erf* Gene Family In Cotton

**DOI:** 10.1101/2020.12.26.423497

**Authors:** Muhammad Mubashar Zafar, Abdul Rehman, Abdul Razzaq, Aqsa Parvaiz, Ghulam Mustafa, Huijuan Mo, Yuan Youlu, Amir Shakeel, Maozhi Ren

## Abstract

*ERF* is a subfamily of *AP2/EREBP* superfamily, contained single AP2 domain. The overexpression of *ERF* genes steered to abiotic stress tolerance and pathogen resistance in transgenic plants. Here, a genome-wide analysis of *ERF* gene family within two diploid species (*G. arboreum & G. raimondii*) and two tetraploid species (*G. barbadense, G. hirsutum*) was performed. A total of 118, 120, 213, 220 genes contained sequence single AP2 domain were identified in *G. arboreum*, *G. raimondii*, *G. barbadense* and *G. hirsutum* respectively. The identified genes were unevenly distributed across 13/26 chromosomes of A and D genomes of cotton. Genome comparison revealed that segmental duplication may have played crucial roles in the expansion of the cotton *ERF* gene family, and tandem duplication also played a minor role. Analysis of RNA-Seq data indicated that cotton *ERF* gene expression levels varied across different tissues and in response to different abiotic stress. Overall, our results could provide valuable information for better understanding the evolution of cotton *ERF* genes and lays a foundation for future investigation in cotton.

## 1. Introduction

Under confrontational environmental conditions such as pathogen attack, submergence, flood, salinity, drought and cold, a precise expression pattern is shown by genes according to their physiological and biological processes [1]. Most importantly, expression of functional genes in the genome is maintained by transcription factors (TFs). During biotic/abiotic stresses and developmental processes, expression of transcription factors of particular genes is enhanced or suppressed by other proteins [2]. Transcription factors for playing vital role as chief regulators in various biological processes are the most important target for crop engineering. [3]. The *AP2/ERF* family is a large group of plant-specific transcription factors. This gene family having *AP2/ERF*-type DNA-binding domain of about 60-70 amino acids was first discovered in *Arabidopsis* homeotic gene, *APETALA2 (AP2)* [4]. *AP2, ERF, RAV* and *DREB* are four subfamilies of *AP2/ERF* gene family. *AP2* subfamily is different from other 3 (*ERF, DREB* and *RAV*) subfamilies as it has double *AP2/ERF* domains while all three have single *AP2/ERF* domain whereas RAV is different from *ERF* and DREB subfamily by having an additional B3 DNA-binding domain. Over the last 20 years, *ERF* family genes caught attention as overexpression of *ERF* genes in different plants directed to abiotic stress tolerance and pathogen resistance in transgenic plants [5, 6].

The *ERF* subfamily members regulate the expression of PR (pathogenesis related) genes through binding to GCC-boxes (AGCCGCC). Further, ERFs are intricated in signaling pathways including salicylic acid, jasmonic acid and ethylene pathways which are important for stress response and plant development [7–9]. Due to their coordinating ability with multiple signaling and hormonal pathways, ERFs are considered excellent entities for engineering biotic/abiotic stress tolerance in plants [9]. Several studies have shown the involvement of *ERF* gene expression under different stress tolerance conditions and tissues in plants [10–12].

Cotton is main agro-industrial crop and is source of most important natural fiber used in textile production [13]. This plays a vital role in global economy and is grown in more than seventy countries. The genus *Gossypium* contains 46 diploid (2n = 2x = 26) and 5 well-established and 1 purported tetraploid (2n = 4x = 52) species. It has been proposed that all diploid cotton species may have evolved from a common ancestor that subsequently diversified to produce eight groups, including groups A–G and K3 [14]. However, biotic and abiotic stresses badly affect the production and growth of cotton. So, struggles to explore the molecular mechanism of stress to support stress tolerance in plants is real and fundamental importance for cotton production [15]. Considering the importance of *ERF* family genes in crop improvement, genome-wide investigation of *ERF* gene-family in cotton can help us to understand the molecular mechanisms of resistance to stress, and thus aid in the development of cotton varieties, using transgenic technology, with greater tolerance to many adverse environments. The release of different cotton whole-genome sequence data, including *Gossypium arboreum* L. [16], *Gossypium raimondii* [17], *Gossypium hirsutum* L. [18] and *Gossypium barbadense* L. [19] has made it possible to systematically identify and analyze the cotton *ERF* genes on a genome-scale level. Here, we performed a comprehensive analysis of cotton *ERF* genes, including their gene structure, motif compositions, chromosome distribution, duplication patterns and expression profiles. This study will provide valuable clues for functional characterization of *ERF* gene family in cotton.

## 2. Materials and methods

### 2.1. Database and sequence retrieval

Gene sequences of *Arabidopsis AtERF* were retrieved from the TAIR (*Arabidopsis thaliana* database) [20]. Cotton genome database Cotton FGD was used for retrieval of *G. barbadense* (NAU), *G. hirsutum* (CRI), *G. raimondii* (JGI) and *G. arboreum* (CRI) genome sequences [21]. *ERF* domain sequence was used as template to retrieve the probable domain homologs from whole genome sequence of *G. barbadense*, *G. hirsutum*, *G. raimondii* and *G. arboreum* [22] through BLASTP at CottonGen (https://www.cottongen.org). All non-redundant hits with less than 1E-5 E-value were taken. Non-targeted and overlapping sequences were removed. Pfam30.0 (http://pfam.xfam.org/) database was used to retrieve hidden Markov Model (HMM) profile of the U-box domain (PF00847) [23], and retrieved results were used as template to find out the candidate *ERF*s from the cotton genome protein database using HMMER3.0 [24]. Protein sequences, CDSs (coding domain sequences) and corresponding full-length sequence in the genome were obtained by using BLAST2.2.31+ (ftp://ftp.ncbi.nlm.nih.gov/blast/executables/blast+/LATEST/). Further, SMART (http://smart.embl-heidelberg.de/) and Pfam 30.0 [23] databases were used for additional analysis to ensure that each candidate protein contained a U-box domain and BUSCA was used for subcellular localization prediction [25, 26].

### 2.2. Gene structure and conserved motif analyses

MEME (Multiple Em for Motif Elicitation) version 5.3.0 (http://meme-suite.org/tools/meme/), an online tool was used for identification of conserved motifs of *ERF* proteins. TBtools software were used to predict gene structure and integrate phylogenetic trees and conserved motifs.

### 2.3. Physical location of ERFs on the chromosome and Phylogenetic tree

Mapcahrt was used to generate the distribution of cotton *ERF* on the chromosome. Genome annotation files were used to retrieve the GFF (general feature format) information of the cotton *ERF*s. The 671 protein sequences of all four species of cotton and 60 protein sequences of *Arabidopsis* were used for the phylogenetic tree. The Clustal Omega (https://www.ebi.ac.uk/Tools/msa/clustalo/; [27] was used to align *ERF* protein sequence and MEGA v7.0 [28] program was used toconstruct a neighbor-joining phylogenetic tree with 1000 bootstrap replicates.

### 2.4. Gene duplication and micro-synteny analysis in *G. arboreum*, *G. raimondii*, *G. barbedense* and *G. hirsutum* L

Information about chromosomal distribution of identified genes were aligned against reference cotton genome in databases. TBtools software were used to visualize the obtained results. Multiple collinear scanning toolkits (MCScanX) and Dual Synteny Plotter software (https://github.com/CJ-Chen/TBtools/) [29] were used for analysis of gene duplication events [30] and syntenic relationship between the *ERF* genes among cotton species, respectively. Some previous studies showed the identification of homologous gene pairs according to MSA (multiple sequence alignment). Advanced circos was used to visualize the collinearity of homologous genes based on the homology between each species and their positions on the genome [29].

### 2.5. Expression analysis

The real expression data were obtained from NCBI database PRJNA171262 for *G. raimondii*, PRJNA594268 for *G. arboreum*, PRJNA490626 for *G. barbadense* and *G. hirsutum* from the sequence read archive (SRA) (https://www.ncbi.nlm.nih.gov) of multiple tissues, flower, seed, organ, ovule, fiber, and under salinity and heat conditions. The Trimmomatic software [31], was used to delete the adapters and to perform quality control. The program hisat2 [32], was used for mapping reads, cufflinks (version: 2.2.1) [33], were used to analyze gene expression levels, and fragments per kilobase million values were used to normalize gene expression levels, then the results were log transformed and a heatmap was generated by MeV [34].

## 3. Results

### 3.1. Identification, Sequence Analysis and Phylogenetic Tree of *ERF* genes in *G. arboreum*, *G. raimondii*, *G. barbadense* and *G. hirsutum*

A total of 118, 120, 213, 220 genes were identified in *G. arboreum*, *G. raimondii*, *G. barbadense* and *G. hirsutum* respectively. To identify *ERF* genes, the sequence of *ERF* domain was blast against the whole genome sequence of *G. arboreum*, *G. raimondii*, *G. barbadense* and *G. hirsutum*. All non-redundant *ERF* genes were obtained from each species. The amino acid sequence of these 671 *ERF* genes were evaluated by Pfam software to confirm their reliability and the presence of *ERF* domains. Genes that lack *ERF* domain in the encoding protein sequences or truncated genes and the genes that were not annotated in their respective genome were deleted. The detailed information of selected genes for *G. arboreum*, *G. raimondii*, *G. hirsutum* and *G. barbadense* are listed in (Supplementary File 1). According to locations on chromosomes the *ERF* genes in cotton were renamed, Gar-ERF-1A.1-13A in *G. arboreum*, Gar-ERF-1D.1-13D in *G. raimondii*, Gba-ERF-1A.1-13D in *G. barbadense*, Ghi-ERF-1A.1-13D in *G. hirsutum*. In *G. barbadense*, 98 *ERF* genes were discovered on At sub-genome, and 115 genes were identified on Dt subgenome. In *G. hirsutum*, 109 genes were identified in At subgenome and 111 genes were identified in Dt subgenome. A phylogenetic (neighbour-joining) tree was constructed to determine the evolutionary relationship of *ERF* genes by using amino acid sequences of identified *ERF* proteins in *G. arboreum*, *G. raimondii*, *G. barbadense*, and *G. hirsutum* with corresponding 60 genes of Arabidopsis (Fig 1). The evolutionary tree classified the *ERF* genes into 8 clades with well supported bootstrap value. The clade one is largest clade followed by clade VI and clade VIII have lowest number of genes followed by clade IV. The results showed that *ERF* genes of four species of cotton and Arabidopsis were unevenly distributed in all clades. Only 2 genes from *G. arboreum*, and 2 from *G. raimondii*, were found in clade VIII.

**Fig 1.**
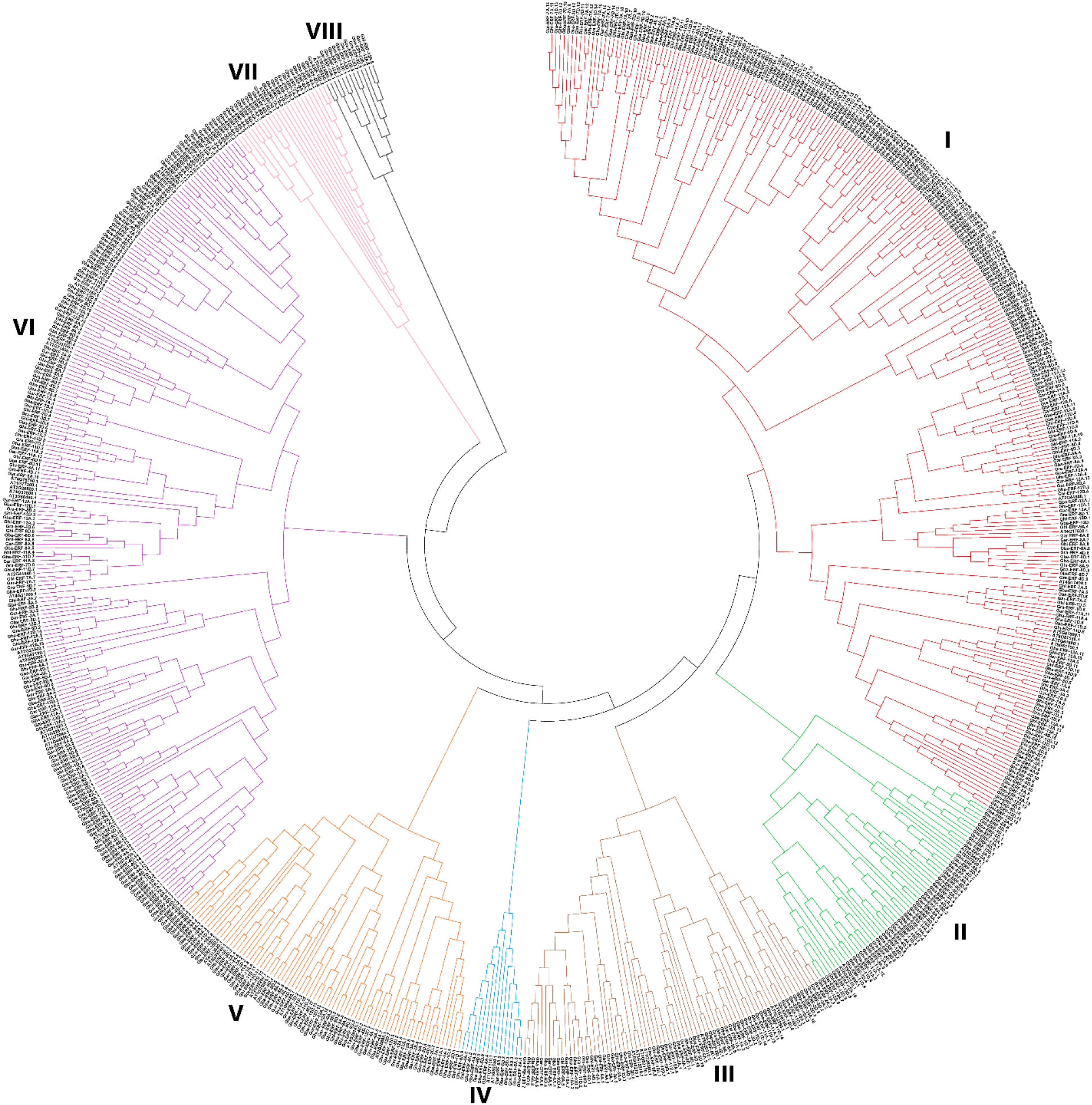
Phylogenetic tree of the *ERF* genes family of cotton and *Arabidopsis*. Bootstrapping values are indicated as percentages along the branches. The different background colorsrs indicate different groups.

### 3.2. Chromosome distribution and gene duplication analysis

To discover the *ERF* genes distribution on chromosomes, each *ERF* gene was mapped on their corresponding chromosome according to gene information of respective genome database. To further examine the evolution of *ERF* genes in four species of cotton, genome duplication events were investigated for WGD or segmental and tandem duplications.

In *G. arboreum*, 118 *ERF* genes were unevenly localized on all 13 chromosomes. The results showed that 7, 5, 9, 2, 9, 8, 14, 14, 8, 8, 15, 15 and 4 genes were located on chromosome A1 to A13 (Fig 2 a). The chromosome 11 and 12 both contained highest number of *ERF* genes (15 ERF) whilst the chromosome 4 and 13 contained lower number of *ERF* genes (2 and 4) respectively. To understand the expansion pattern of *ERF* gene family in *G. arboreum*, the circos analysis was performed. The results revealed that 92 *ERF* genes have WGD or segmental duplication and located on all chromosomes (Fig 3 A). The 13 *ERF* family genes were tandemly duplicated and distributed on chromosomes 2, 5, 7, 8, 9 and 12. Other 11 genes were dispersed in *G. arboreum* genome (Supplementary File 2).

**Fig 2 (a, b).**
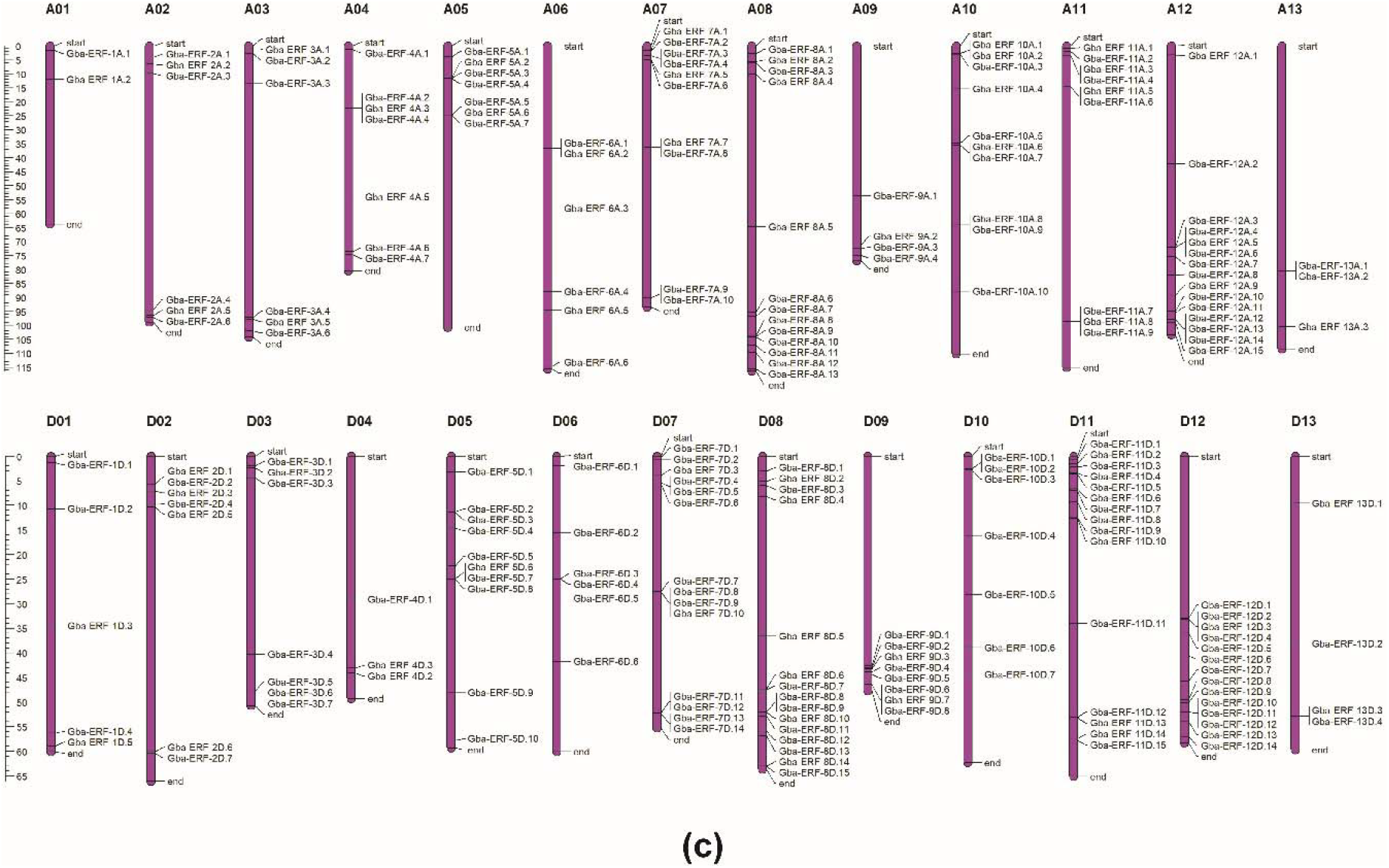
Gene Location on chromosome of *G. arboreum* and *G. raimondii.*

In *G. raimondii*, 120 *ERF* genes were distributed unequally on all 13 chromosomes (Fig 2 b). The number of *ERF* genes from chromosome D01 to D13 were 14, 5, 8, 15, 8, 8, 15, 17, 11, 7, 7, 2 and 3 respectively. The chromosome D08 and D04 contain highest number of genes 17 and 15 respectively. While the chromosome D12 and D13 exhibits lowest number of genes 2 and 3 respectively. Among120 *ERF* genes 74 genes have WGD or segmental duplications and located unevenly on all chromosomes (Fig 3 B). The chromosomes D08 and D01 contain highest WGD or segmentally duplicated genes 14 and 10 respectively. On chromosome D12 and 13 only single gene have segmental duplications (Fig 3 B). The genes Gra-ERF-1D.6, Gra-ERF-1D.13, Gra-ERF-4D.3, Gra-ERF-4D.9, Gra-ERF-6D.6, Gra-ERF-6D.7, Gra-ERF-6D.8, Gra-ERF-7D.14, Gra-ERF-8D.15, Gra-ERF-9D.4, Gra-ERF-9D.7, Gra-ERF-9D.8, and Gra-ERF-13D.3 have tandem duplications (Supplementary File 2).

In *G. barbadense*, 213 *ERF* genes were scattered on all 26 chromosomes of At and Dt subgenomes. The 98 and 115 *ERF* genes were identified in At and Dt subgenomes respectively (Fig 2 C). In At genome of *G. barbadense*, the chromosome A12 had maximum (15) *ERF* genes whilst the chromosome A01 and A13 have minimum number of *ERF* genes 2 and 3 respectively. In Dt subgenome the chromosome D08 and D11 contains greatest number of (15) *ERF* genes and chromosome D04 had lowest (3) *ERF* genes. The 182 *ERF* genes of *G. barbadense*, have segmental duplications and these were scattered unevenly on all 26 chromosomes. Out of these 182 genes 42.75% and 57.14% were belongs to At and Dt subgenome respectively (Fig 3C). The chromosome A13 have only single genes Gba-ERF-13A.3 had segmental duplication. From At subgenome, the genes Gba-ERF-4A.3, Gba-ERF-4A.4, Gba-ERF-5A.6, Gba-ERF-6A.2, Gba-ERF-7A.4, Gba-ERF-7A.9, Gba-ERF-8A.4, Gba-ERF-10A.7, Gba-ERF-11A.8, Gba-ERF-11A.9, Gba-ERF-13A.1 and Gba-ERF-13A.2 have tandem duplications (Supplementary File 2). Only three genes Gba-ERF-7D.13, Gba-ERF-7D.14, and Gba-ERF-11D.13 from Dt subgenome have tandem duplications (Supplementary File 2).

**Fig 2 c.**
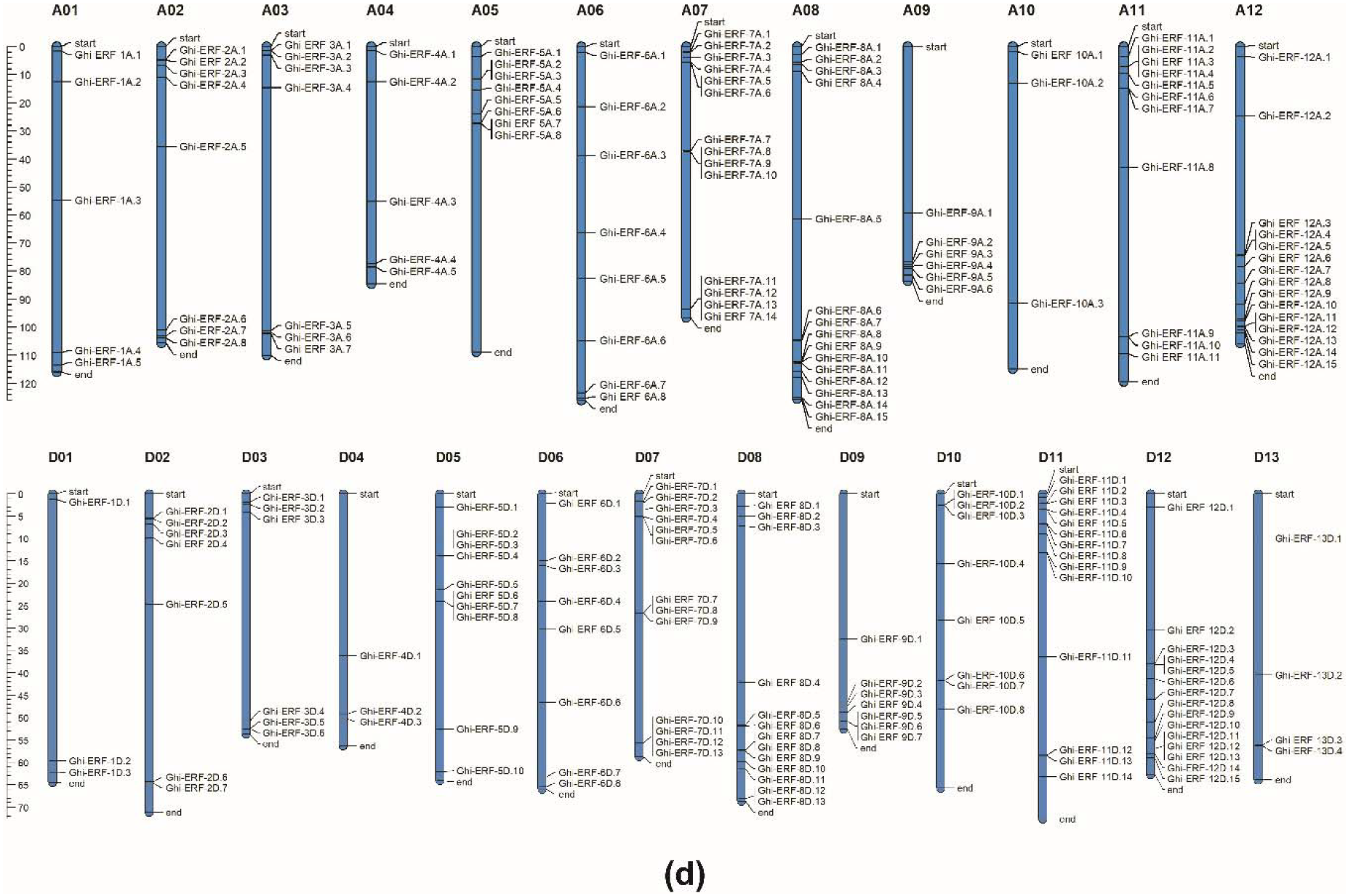
Gene Location on chromosome of *G. barbadense*.

In *G. hirsutum*, 220 *ERF* genes were unevenly distributed on all 26 chromosomes. Among these 220, 109 and 111 *ERF* genes were located in At and Dt subgenome respectively (Fig 2D). In At genome of *G. hirsutum*, the chromosome A08 and A12 have maximum (15) *ERF* genes whilst the chromosome A10 and A13 have minimum number of *ERF* genes 3 and 4 respectively. In Dt subgenome the chromosome D12 contains greatest number of (15) *ERF* genes and chromosome D01 and D04 have lowest (3) *ERF* genes (Fig 2D). The segmental duplications were found in 94.55% *ERF* genes of *G. hirsutum*, 49.52% and 50.48% genes were located in At and Dt subgenomes respectively (Fig 3D and Supplementary File 2). Tandem duplications were found in 3.18% *ERF* genes and these genes were located on chromosome A07, A08, D07, D08, D09 and D11.

**Fig 2 d.**
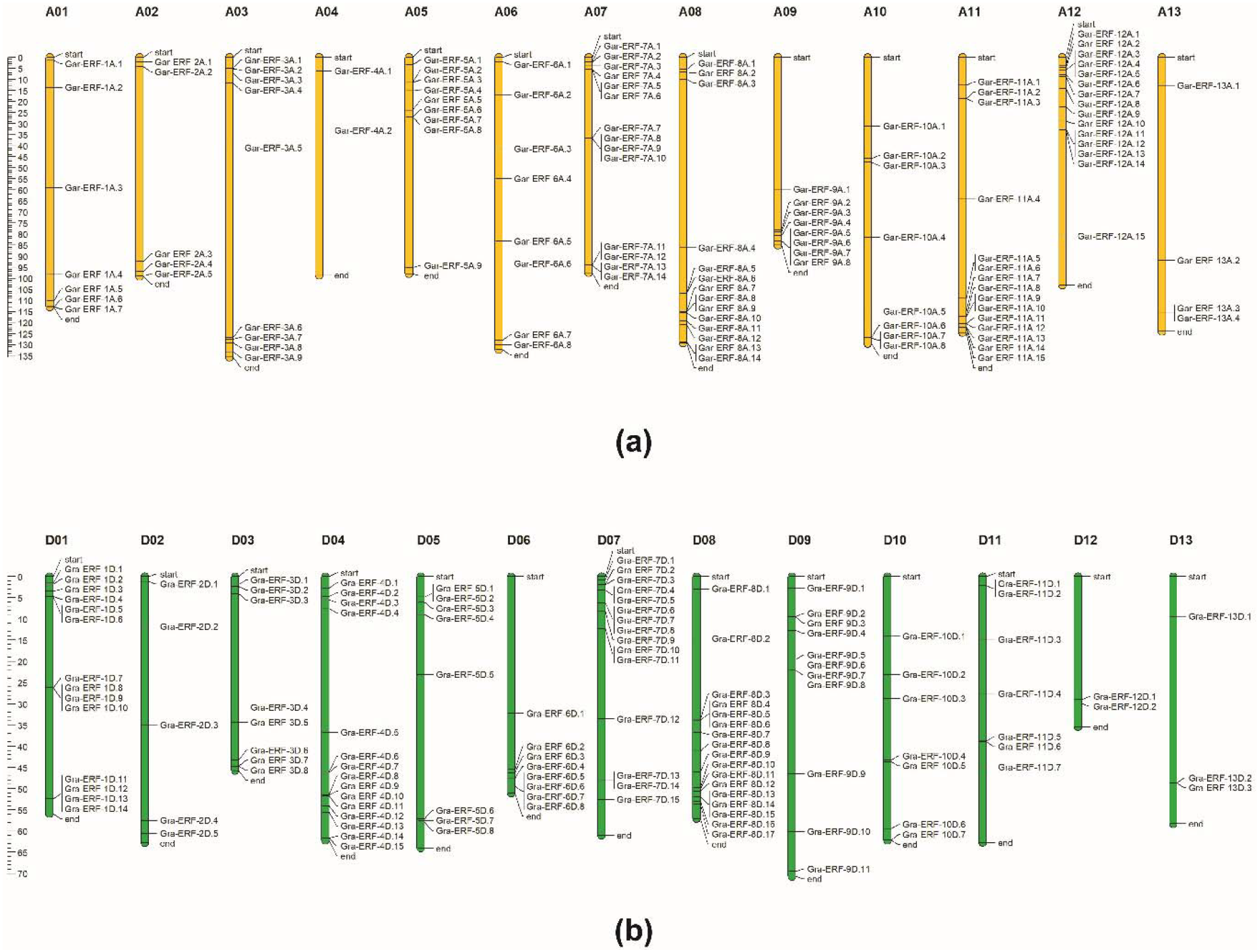
Gene Location on chromosome of *G. hirsutum*.

**Fig 3.**
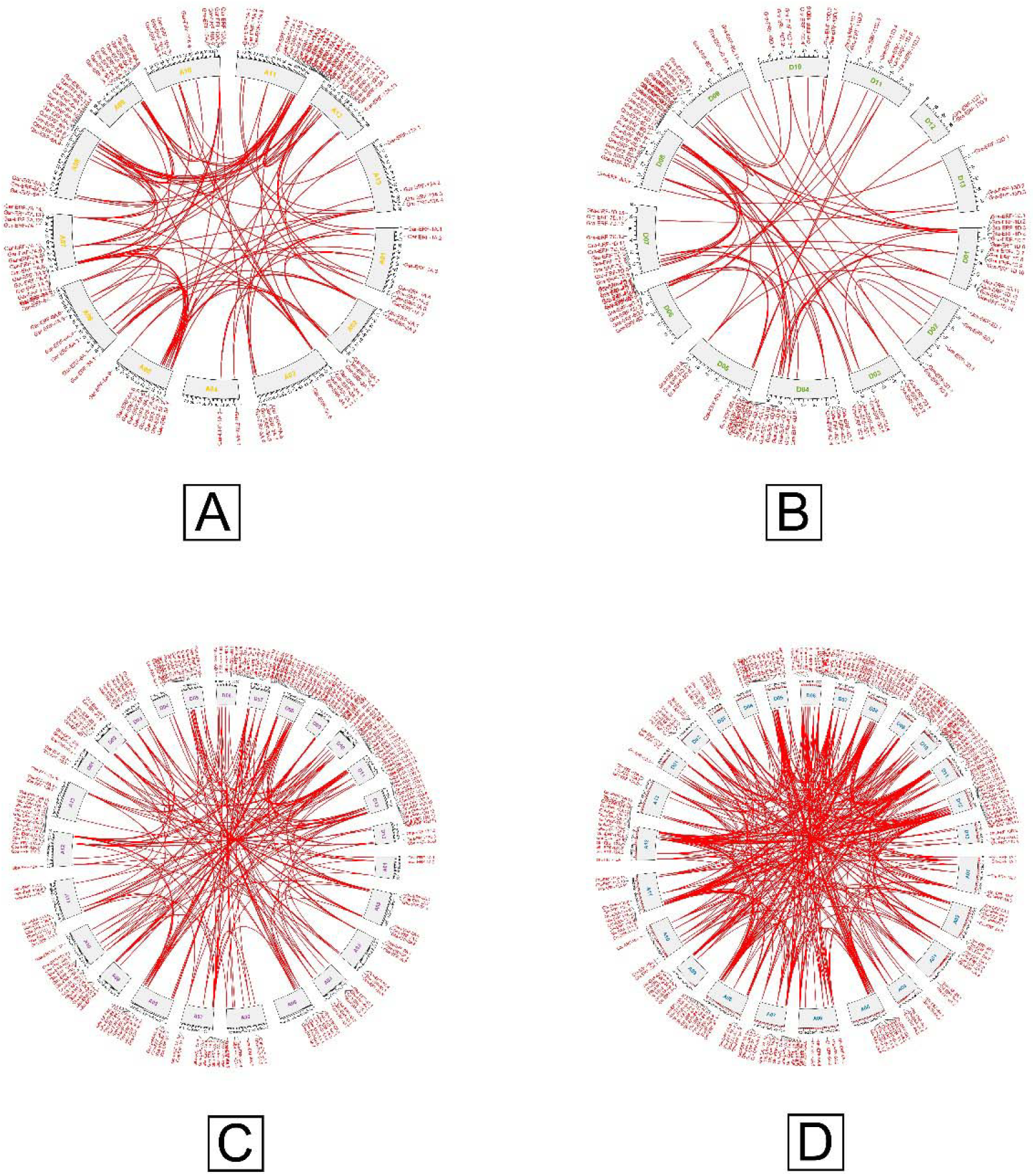
Duplicated *ERF* gene pairs identified in *G. arboreum, G. raimondii, G. barbadense* and *G. hirsutum.*

The *G. hirsutum* and *G. barbadense* are evolved due to hybridization between an A-genome species (G. herbaceum or *G. arboreum*) and a D-genome genome (*G. raimondii*) [35]. To understand the evolutionary relationships of *ERF* genes, a relative syntenic map of *ERF* genes from the four cotton species was fabricated (Fig 4). Synteny analysis showed several gene loci that are highly conserved between the At and Dt sub-genomes of both tetraploid cotton species. According to our MCScan analysis, 237, 105 duplication gene-pairs were found between diploid *G. arboreum* and tetraploid *G. barbadense*, *G. hirsutum* respectively. 393, 307 duplication gene-pairs were found between diploid *G. raimondii* and tetraploid *G. hirsutum*, *G. barbadense* respectively. The location of *ERF* genes on D11, D01, D04 and D06 in *G. raimondii* have good collinear relationship with the *ERF* genes present on the homologous chromosomes of *G. hirsutum* and *G. barbadense* (Fig 4). The *ERF* genes on A11 in *G. arboreum* also have good collinear relationship with the *ERF* genes present on the homologous chromosomes of *G. hirsutum* and *G. barbadense*. The synteny map revealed that small deletion, duplication and reshuffling of chromosome may have occurred during evolution.

**Fig 4.**
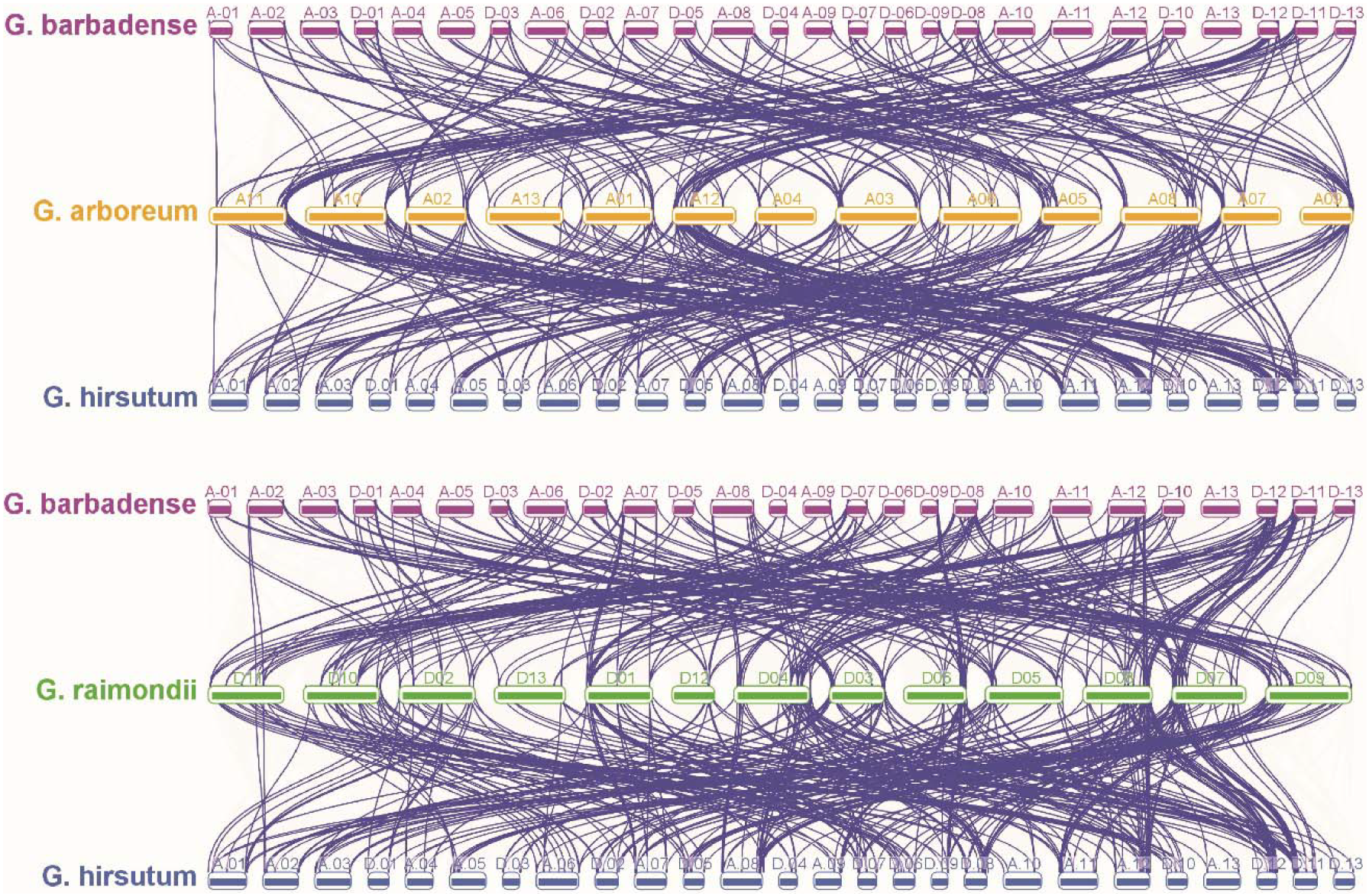
The sub-genome distribution and synteny analysis of cotton *ERF* genes. The blue lines indicate duplicated *ERF* pairs, the gray lines indicate collinear blocks.

### 3.3. Gene structure and conserved motifs of the cotton ERF gene family

For more understanding into the evolution and structural diversity of the *ERF* family in each cotton species, we analyzed the gene structure and conserved motif of the *ERF* genes. The motifs logo, motifs sequences, motifs name, motif width and motif similarity matrics in 4 species of cotton are provided in (Supplementary File 3). The motif 1 and 2 were conserved in all *ERF* genes across all four species. In *G. arboreum*, based on evolutionary tree the *ERF* family genes are categorized in 3 groups. As shown in (Fig 5 A) only 16 genes have introns and accounting for 13.56 % and remaining 86.44 % genes are intronless. All the *ERF* family genes have conserved exon number (1) except the 16 genes that contain introns have 2 exon number. The transcript length of 118 *ERF* genes of *G. arboreum*, are 384-1269 bp, and the protein sequences are 127-422 aa. The Isoelectric Point of these proteins exist between 4.292 and 10.542, and the molecular weight is between 14.323 kDa and 47.137 kDa (Supplementary File 1). To gain more insight about divergence and functional relationship of Gar-ERF proteins, a total of 12 conserved motifs in the cotton *ERF* were recognized by MEME software, and the height of each letter in the logo was proportional to the conservation level of amino acid in all sequences analyzed. As shown in (Fig 5 A) all the Gar-ERF proteins contain motif 1 and 2. The motifs in different groups indicated that varying degrees of divergence among them. The genes Gar-ERF-11A.12 and Gar-ERF-12A.7 were distinct in group 1 due to presence of motif 12. The presence of motif 3 in some genes of group 2 and 3 make them different from other genes. In general, the *G. arboreum*, *ERF* proteins in the same group usually contained similar motifs, which indicates that they may play similar roles in the development and growth of *G. arboreum*.

**Fig 5A.**
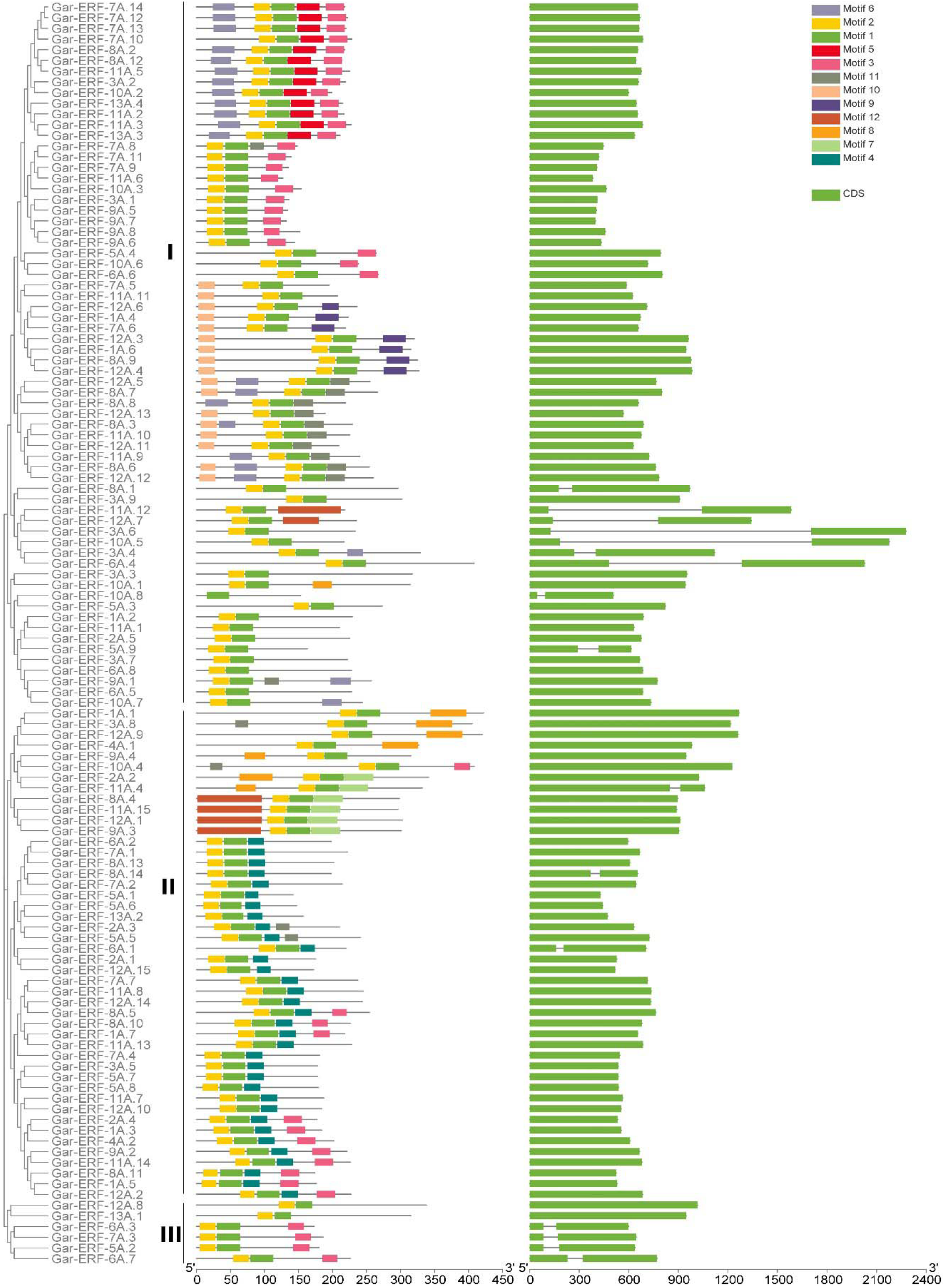
Evolutionary tree with motif and gene architecture of *ERF* gene family *in G. arboreum*.

The transcript length of 120 *ERF* genes of *G. raimondii* are 399-2134 bp, and the protein sequences are 127-420 aa. The Isoelectric Point of these proteins exist between 4.312 and 10.846, and the molecular weight is between 14.318 kDa and 47.139 kDa. The 80.83 % genes were intron-less and 19.16 % genes have introns (Fig 5 B & Supplementary File 1). The gene Gra-ERF-1D.9 had highest mean intron length 3401 bp. The 81.66 %, 15 %, and 4.16 % genes have exon number 1, 2 and 3 respectively. The mean exon length are ranges between 260-2134 bp. The motif 3, 4, 5, 6, 7, 8, 9, 10, 11 and 12 were detected in 40, 30, 13, 20, 5, 18, 7, 5, 5, and 8 genes respectively. The genes were divided in 3 groups on the bases of evolutionary tree. In group 1 the genes Gra-ERF-3D.8, Gra-ERF-8D.15, Gra-ERF-4D.10, Gra-ERF-8D.6, Gra-ERF-8D.14 were distinct due to presence of motif 10. In group 2, only one gene was unique due to presence of motif 3 (Fig 5 B).

**Fig 5B.**
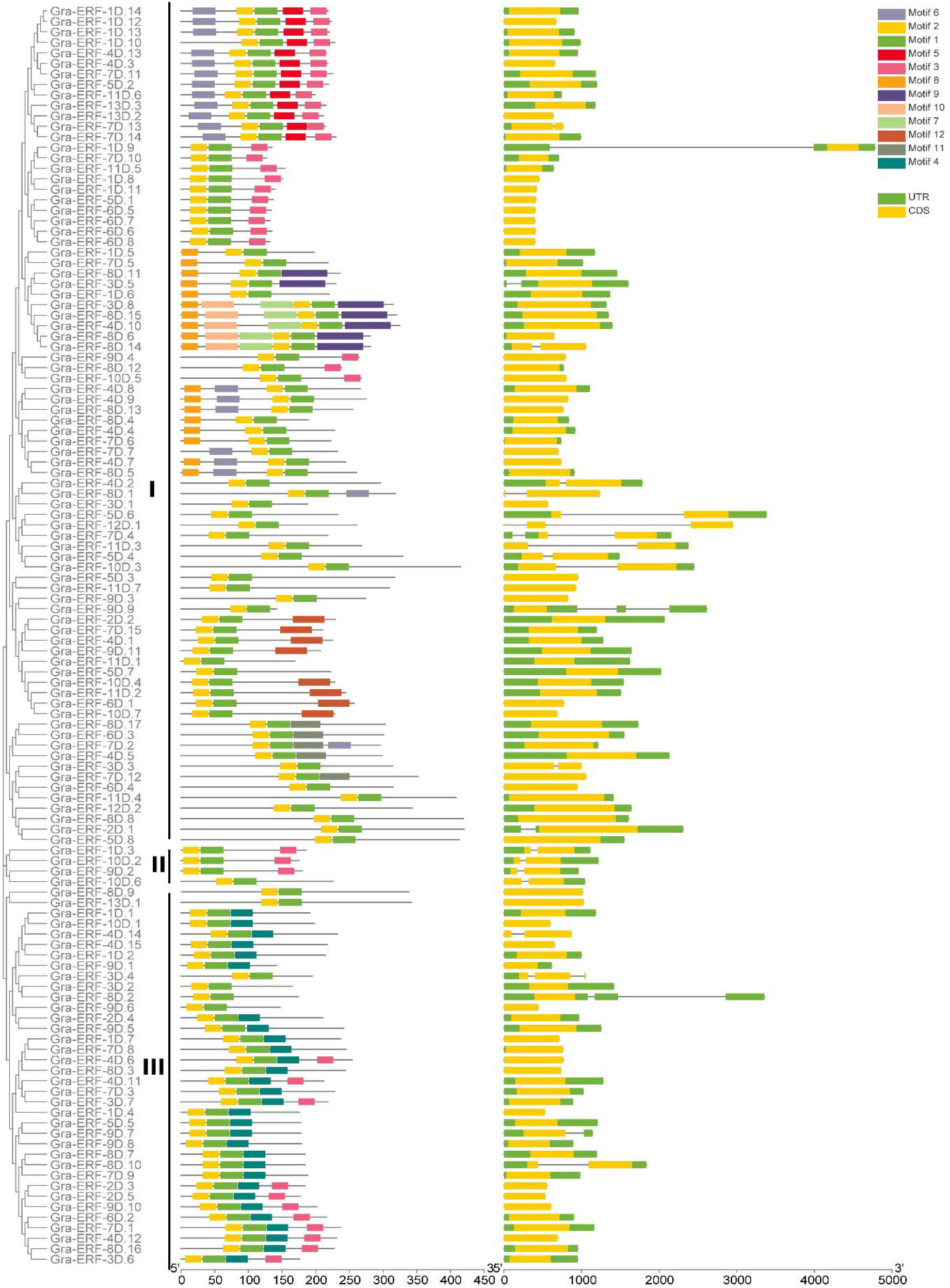
Evolutionary tree with motif and gene architecture of *ERF* gene family *in G. raimondii.*

The transcript length of 213 *ERF* genes of *G. barbadense*, are 339-1914 bp. The protein sequences are ranges between 112-637 aa. The Isoelectric Point of these proteins exist between 4.312 and 11.24, and the molecular weight is between 12.866 kDa and 71.803 kDa. The 62.44 % genes were intron-less and 37.55 % genes have introns. The gene Gba-ERF-4A.6 had highest mean intron length 3145 bp while the gene Gba-ERF-3D.1 lowest mean intron length 27 bp. The exon number are 1-4 in all *ERF* genes of *G. barbadense*, and mean exon length are 119-1275 (Supplementary File 1). The motif 3, 4, 5, 6, 7, 8, 9, 10, 11 and 12 were detected in 25, 39, 45, 35, 49, 14, 13, 17, 41 and 19 genes respectively (Fig 5 C).

**Fig 5C.**
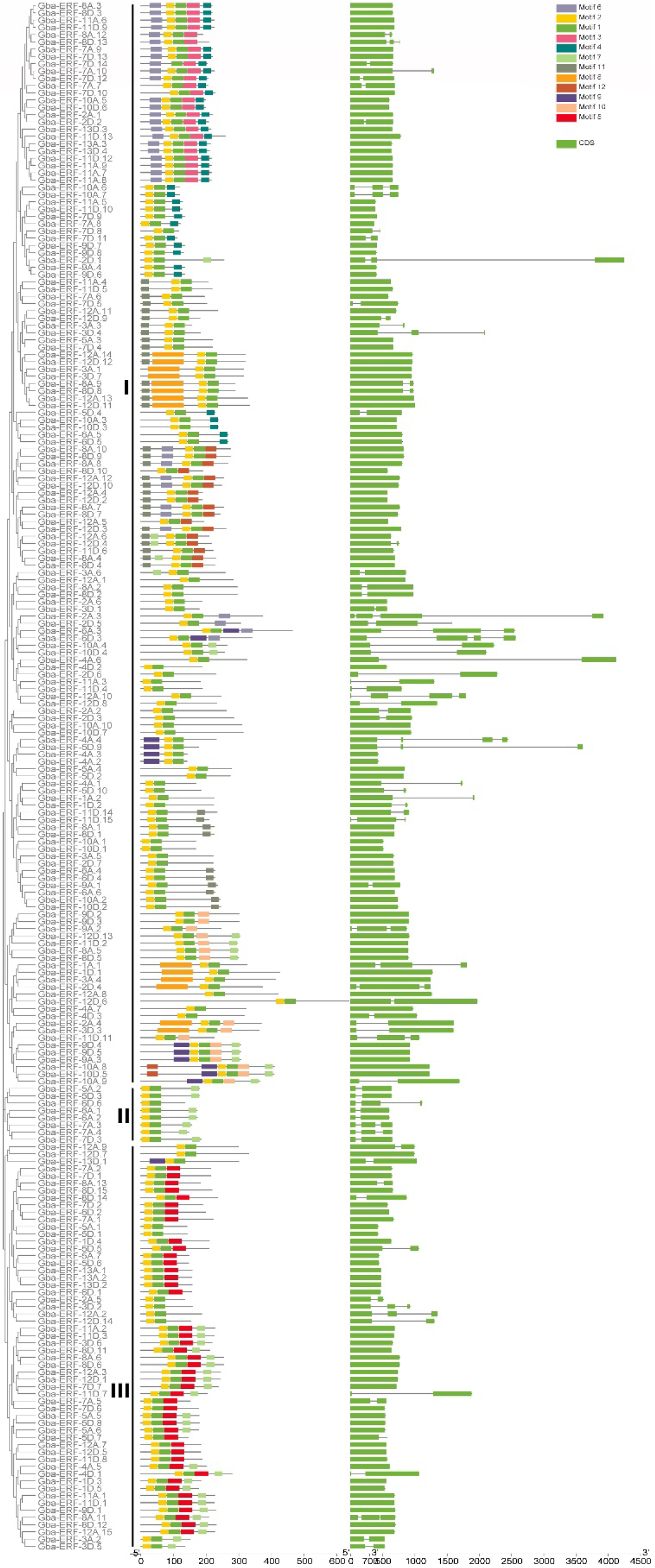
Evolutionary tree with motif and gene architecture of *ERF* gene family *in G. barbadense.*

The transcript length of 220 *ERF* genes of *G. hirsutum*, are 327-10114 bp. The protein sequences are ranges between 108-483 aa. The Isoelectric Point of these proteins exist between 4.284 and 10.846, and the molecular weight is between 11.992 kDa and 52.887 kDa. The 86.36 % genes were intron-less and 13.64 % genes have introns. The gene Ghi-ERF-10A.2 had highest mean intron length 2202 bp while the genes Ghi-ERF-5A.1 and Ghi-ERF-3D.3 have least intron length 27 bp. Most of the genes 190, and 27 genes have 1 and 2 exon number (Supplementary File 1). The mean exon length is 119.5-1839. The motif 3, 4, 5, 6, 7, 8, 9, 10, 11 and 12 were detected in 60, 80, 25, 32, 16, 11, 15, 12, 26 and 8 genes respectively (Fig 5 D).

**Fig 5D.**
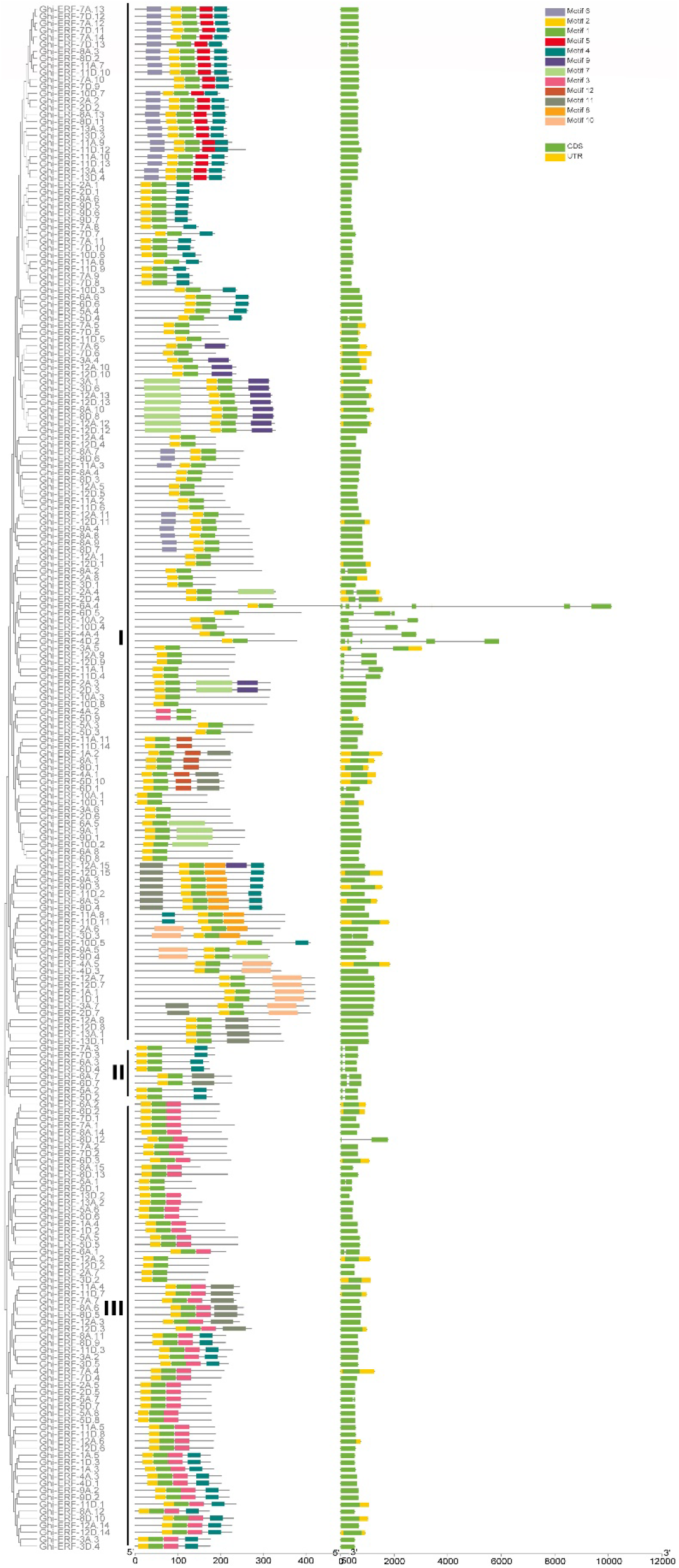
Evolutionary tree with motif and gene architecture of *ERF* gene family *in G. hirsutum.*

### 3.4. Subcellular localization

The 93.22 % *ERF* genes of *G. arboreum*, were located in nucleus. The genes Gar-ERF-3A.9, Gar-ERF-5A.2, Gar-ERF-6A.3 and Gar-ERF-8A.10 were found in chloroplast. Only 3 genes namely Gar-ERF-6A.7, Gar-ERF-8A.3 and Gar-ERF-8A.6 were located in extra cellular spaces. Only Gar-ERF-1A.2 gene was present in mitochondria (Supplementary File 1). The 92.5 % and 4.16 % *ERF* genes of *G. raimondii*, were located in nucleus and chloroplast respectively (Supplementary File 1). Only 1 and 3 genes were located in mitochondria and extracellular spaces respectively. The 89.20 % and 5.63 % *ERF* genes of *G. barbadense*, were located in nucleus and chloroplast respectively (Supplementary File 1). Only 5 genes were located in extracellular spaces. The 91.81 %, 4.54 % and 2.73 %, *ERF* genes of *G. hirsutum*, were located in nucleus, chloroplast and extracellular spaces, respectively (Supplementary File 1). Only Ghi-ERF-1A.2 and Ghi-ERF-4A.4 genes were located in mitochondria and plasma membrane respectively.

### 3.5. Expression analysis

Different cotton *ERF* genes across different tissues like seed, stem, leaves, flower and fiber. RNA-seq data for fiber (0 DPA ovules, 1 DPA ovules, 3 DPA ovules, 5 DPA and 10 DPA fibers) and seed (10 DPA, 20 DPA, 30 DPA and 40 DPA) of *G. arboreum*, was used for expression analysis. The Gar-ERF genes were clustered in 5 pattern groups on the basis of heat map of fiber and seed. Generally, similar expression pattern was observed within groups. The genes of group 1, 2, 3, 4 and 5 were highly expressed in ovule 5DPA, 3DPA, 1DPA, 0DPA and fiber 10DPA respectively. From group first only Gar-ERF-4A.1, Gar-ERF-5A.1 and Gar-ERF-11A.7 were expressed in fiber 10DPA. Some genes from group 5 were also expressed in ovule 0DPA (Fig 6 a).

**Fig 6A.**
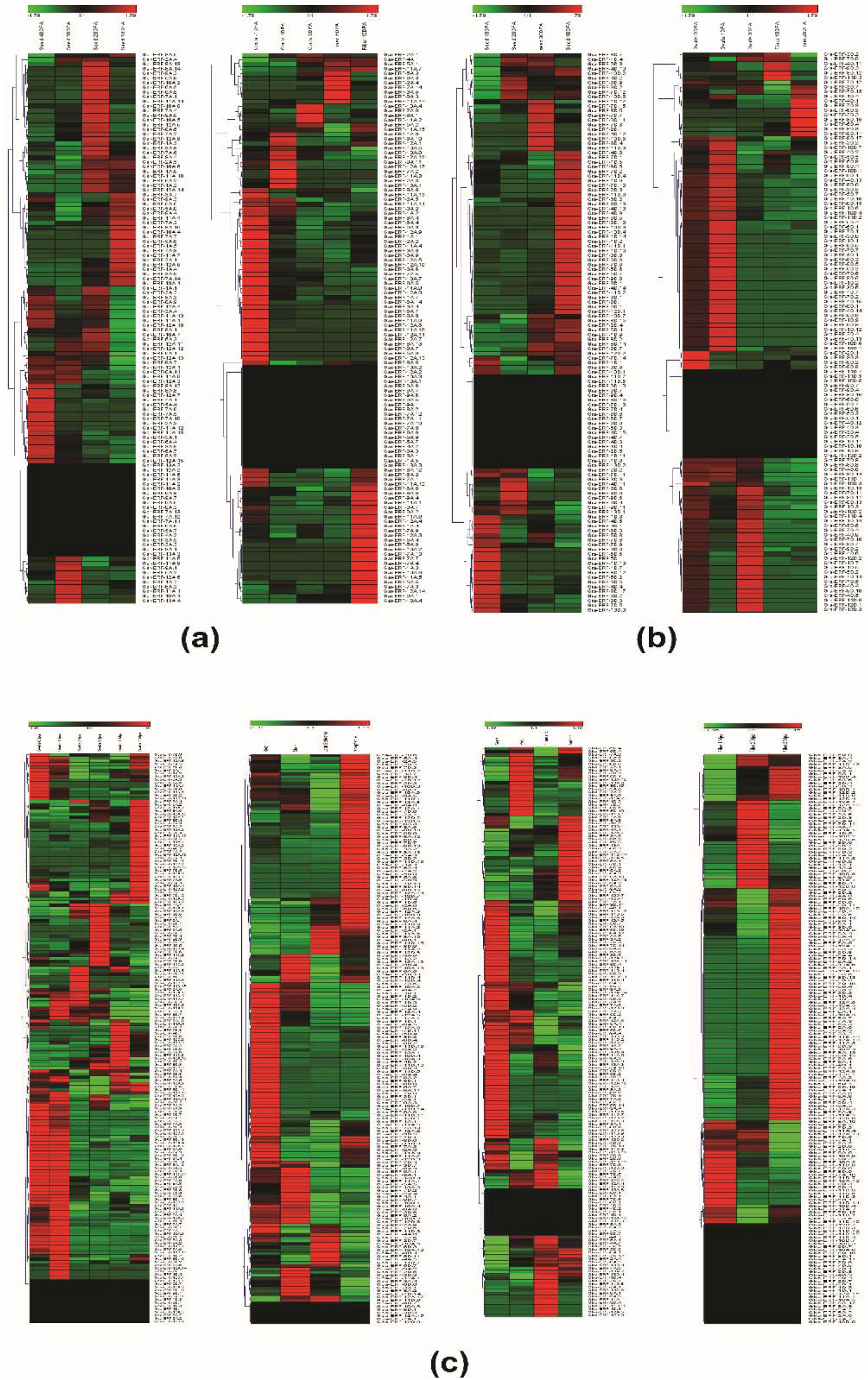
Expression patterns of *ERF* genes (Fig a & b) in seed, ovule and fiber of *G. arboreum,* and *G. raimondii* and in (Fig c) expression patterns of *ERF* genes in seed, ovule fiber, tissue and flower of *G. barbadense*.

In case of *G. raimondii*, the genes of group 1, 2, 4 and 5 were highly expressed in seed 20 DPA, 10 DPA, 40 DPA and 30 DPA respectively. The genes of group 3 were highly expressed in seed at 40 DPA and 30 DPA. In case of seed, heat map of Gra-ERF genes were clustered in six pattern groups (Fig 6 b). Most of the genes of group 1, 2, 3, 4, 5 and 6 were highly expressed in seed at 20 DPA, 30 DPA, 40 DPA, 30 DPA, 20DPA and 10 DPA respectively. The genes of group 1, 2, 3, 4 and 5 were highly expressed in fiber 10DPA, fiber 20DPA, ovule 1DPA, 0DPA and 3DPA respectively (Fig 6 b).

**Fig 6B.**
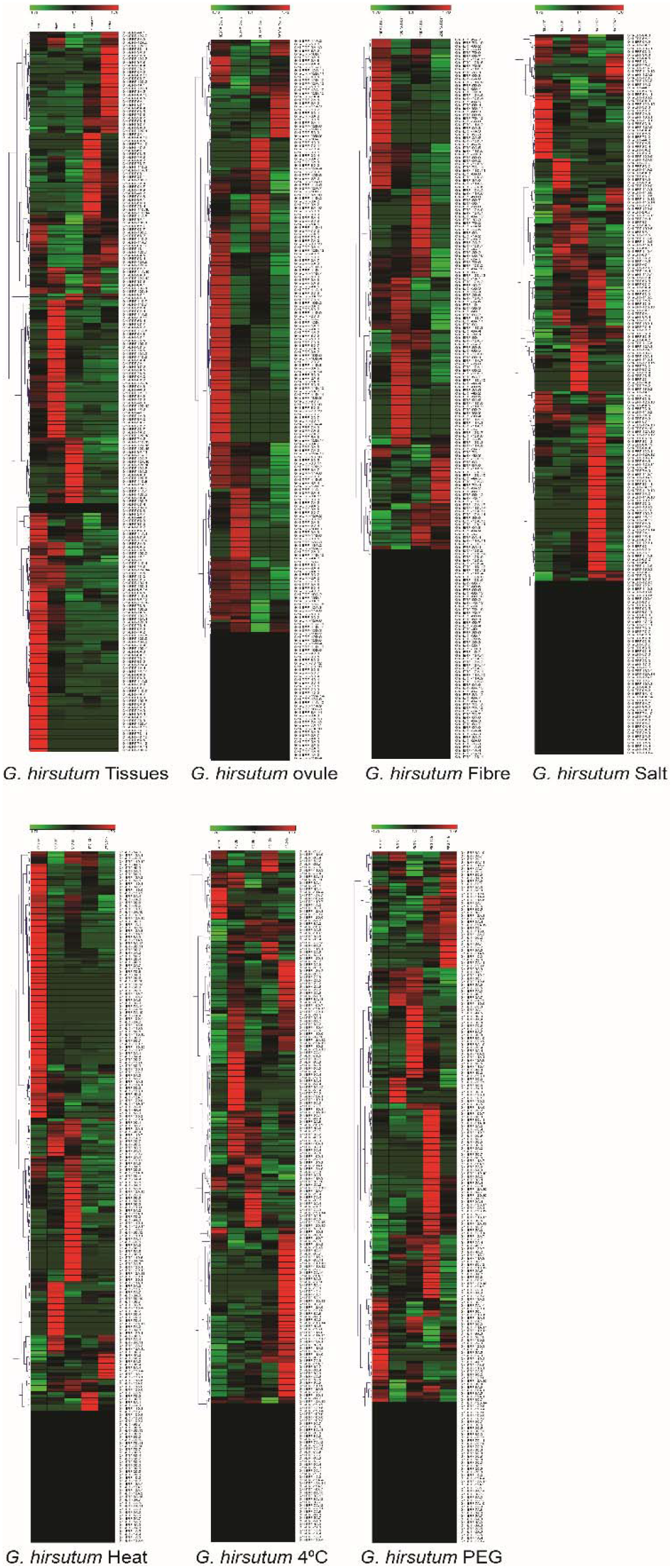
Expression patterns of *ERF* genes in tissue, ovule, fiber and (salinity, heat, cold and PEG) conditions at for (1h, 3h, 6h, 12h and 24h) in *G. hirsutum.*

The expression profile of Gba-ERF genes in ovule (0DPA, 1DPA, 3DPA, 5DPA, 10DPA, 20DPA) organs (root, stem, leaf apicalyx, receptacles), flower (sepal, petal, filament, anther) and fiber (10 DPA ovules, 20 DPA ovules, 25 DPA ovules) were investigated. The expression pattern of Gba-ERF genes in ovule clustered the genes in 5 groups (Fig 6 c). The genes of group 1, and 2 were expressed in ovule at 0DPA, and 20DPA respectively. The genes of group 3 were mostly expressed in 5DPA followed by 3DPA and 1DPA respectively. Most of the genes of group 4 were expressed at 10DPA. The genes of group 5 were highly expressed in 0DPA and 1DPA. The heat map of Gba-ERF genes in organ clustered the genes into 4 groups on bases of their expression pattern (Fig 6 c). The genes of 1^st^ group were highly expressed in receptacle and few genes of 1^st^ group in leaf apicalyx. Most of the genes of group 2 were highly expressed in root. The genes of group 3 and 4 were expressed in stem and leaf apicalyx respectively. In case of flower and fiber the heat map also clustered the genes into 4 groups. In flower the genes of group 1, 2, 3 and 4 were highly expressed in petal, anther, sepal and filament respectively (Fig 6 c). Most of the genes of group 1 and 3 were highly expressed in fiber at 25DPA but the genes Gba-ERF-5D.8, Gba-ERF-3A.4, Gba-ERF-7A.2, Gba-ERF-11D.13 from 1^st^ group were expressed in fiber at 20 DPA. The genes of group 2 were expressed in fiber at 20 DPA. The genes of group 4 were highly expressed in fiber at 10 DPA (Fig6 c).

In *G. hirsutum*, the Ghi-ERF genes expression was examined in organs (root, stem, leaf, filaments, anther), ovule (10DPA, 15DPA, 20DPA, 25DPA), fiber (10DPA, 15DPA, 20DPA, 25DPA), salinity, heat, cold, and PEG stress for (1hour, 3hours, 6hours, 12hours and 24hours) respectively. On the basis of Ghi-ERF genes expression in organs (root, stem, leaf, filaments and anther), the genes were clustered in 5 groups (Fig 6 d). Most of the genes of 1, 2, 3, 4 and 5 were highly expressed in anther, filaments, stem, leaf and root respectively. In ovule and fiber, based on heat map Ghi-ERF genes were clustered in 4 expression pattern groups (Fig 6 d). For ovule, most of the genes of group 1, 2, 3 and 4 were highly expressed at 25DPA, 20DPA, 10DPA, and 15DPA respectively. Most of the genes of group 1, 2, 3 and 4 were highly expressed in fiber at 10DPA, 20DPA, 15DPA, and 25DPA respectively. Under salt stress, heat map clustered the genes in 3 groups (Fig 6 d). Most of the Ghi-ERF genes of group 1 were highly expressed at NaCl 1hour, while some genes of group 1 were also expressed at NaCl 24 hour. The Ghi-ERF genes of 2^nd^ group were differentially expressed at NaCl 3hours, NaCl 6hours, and NaCl 12hours. Most of the genes of group 3 were highly expressed at NaCl 12hours. The heat map of Ghi-ERF genes in heat stress divide the genes in three groups. The genes of group 1 were highly expressed at 37 C 1hour. Most of the genes of 2^nd^ and 3^rd^ group were highly expressed at 37 C 6hours and 37 C 3hours respectively (Fig 6 d). Some genes of group 3 were also expressed at 37 C in 24h and 12 h. The expression of *ERF* genes under cold and PEG stress the heat map clustered these genes in 5 groups. In case of cold stress most of the genes of group 1 were expressed at 4C in 1h and 12h. The genes of group 2, 3, 4 and 5 were highly expressed at 4C in 24h, 3h, 6h and 24h respectively. In case of PEG stress, most of the genes of group 1, 2, 3, 4 and 5 were highly expressed in 24h, 6h, 3h, 12h and 1h respectively (Fig 6 d).

## 4. Discussion

In plants, the (*AP2/EREBP*) superfamily is one of the important and largest transcription factor (TF) families. *AP2/ERF* domain specifies different members of the (*AP2/EREBP*) family. The domain consists of up to 60-70 amino acids [4]. These members have necessary roles in the growth and development of plants, and in responding a variety of environmental stresses involving heat, drought, salinity, cold and various pathogen infections. The roles are exerted either by direct response to the stresses or by regulation of expression of target genes downstream [36]. Numerous studies have recognized different numbers of *ERF* family members in plants; like in soybean 120 *ERF* family members [37], cucumber 103 (*Hu & Liu, 2011*) tomato 85 [38], rice 139 [39] and 147 Arabidopsis [40] respectively.

Arabidopsis, used as the model plant comprise of 147 *AP2/EREBP* genes divided into 4 subfamilies which are; APETALA2 (AP2), Ethylene-Responsive Factor (ERF), Dehydration-Responsive Element Binding protein (DREB), and Related to ABI3/VP1 (RAV) subfamilies [41]. Subfamily members of *ERF* and DERB containing single conserved domain AP2 and making the largest groups in superfamily *AP2/EREBP*, perform important roles in various plant processes and variety of stress responses.The sequences containing one AP2 domain have biggest number of members in the superfamily *AP2/EREBP*, which have been further divided into 2 major subfamilies; *ERF* subfamily and DREB subfamily [42].

Cotton is an important economic crop in the world providing the leading share of world’s natural textile fiber and a substantial amount of edible oil. Studies revealed the formation of the allotetraploid Gossypium species occurred about 1-1.5 million years back through a polyploidization event which involved two species; one being A-genome species called *G. arboreum* and the other was paternal D-genome species named *G. raimondii* [18].

WGD or polyploidy is being acknowledged as a source of evolution in plants, a significant force resulting in enormous silencing and removal of duplicated genes [43]. In this study, we identified 118, 120, 213, 220 *ERF* genes in *G. arboreum*, *G. raimondii*, *G. barbadense* and *G. hirsutum* respectively. The evolutionary tree showed that more *ERF* proteins appeared in pairs and clustered together with 2 Gba-ERF, 2 Ghi-ERF, 1 Gar-ERF and 1 Gra-ERF which supported the cotton species polyploidization event that occurred 1.5 million years ago [44].

The tetraploid cotton should contain *ERF* genes in a number equal to *ERF* genes contained in *G. arboreum* plus *ERF* genes in *G. raimondii*. Though the number of *ERF* genes actually in allotetraploids (*G. barbadense* 213 and *G. hirsutum* 220) appeared lesser than those in two diploids (*G. arboreum* 118 and *G. raimondii* 120), implying that the chromosomes doubling and then genome rapid sequence arrangement would produce gene loss of different degrees in polyploidization process [18]. Gene duplication is a significant source of evolution in genome and in turn genetic system by producing new gene subfamilies [45]. Polyploidy, segmental and Tandem duplications chiefly help to generate new gene families [45]. *ERF* genes in *G. arboreum*, (13), *G. barbadense* (15), *G. hirsutum* (7) and *G. raimondii*, (13) have tandem duplications. Segmental duplications frequently take place in plants as most plants occur as diploidized polyploids [46]. In *G. arboreum* (92), *G. hirsutum* (208), *G. barbadense* (182) and *G. raimondii* (74) *ERF* genes have undergone to segmental or WGD duplications.

There’s a possibility of deletion of some of pre-existing genes or newly generated genes during the process of cotton evolution. Collinearity and chromosomal locations showed the significant role of segmental duplications in *ERF* genes expansion in cotton which was in accordance with the results reported in rice, B. napus, A. thaliana and other plants [10, 38, 47, 48]. The synteny analysis results could be used to reveal the evolutional and functional connections among the cotton species. In these studies, 237, 105 duplication gene-pairs were established between diploid *G. arboreum* and tetraploid *G. barbadense*, *G. hirsutum* correspondingly. And there were 393, 307 duplication gene-pairs established between diploid *G. raimondii* and tetraploid *G. hirsutum*, *G. barbadense* correspondingly. Our study analysis showed a higher homology between *ERF* genes of tetraploid and diploid cotton species. Some *ERF* genes did not find any orthologous gene pairs, which could be attributed to chromosomes rearrangement or fusion during the evolution [49, 50].

Gene structure analysis plays a crucial role in revealing the function of genes [47]. Here, our results suggested that 13.56 %, 19.16 %, 37.55 % and 13.64 % genes possess introns in *G. arboreum*, *G. raimondii*, *G. barbadense* and *G. hirsutum* respectively. The *ERF* members demonstrated similar gene structure within same group. Loss of introns occurred more rapidly in genes than intron acquisition after segmental duplication [51]. Also, some studies revealed that intron number and distribution were related to evolution in plant [52] in a way that introns possibly have been lost during evolution from *ERF* family genes in higher plants. Our results showed that 86.44 %, 80.83 %, 62.44 %, and 86.36 % *ERF* genes have no introns which is similar to the status in *Arabidopsis*, cucumber and rice [53–55]. Transcriptional output can be delayed due to long multiple introns, possibly causing suppression of genes expression in adverse conditions. Contrarily, the genes containing small or fewer introns might have efficient expression while responding to stress environments [56]. Therefore many of intron-less *ERF* genes might react quickly to the external environment variations [47]. The transcription factors domains and motifs perform essential roles during transcriptional activity, proteins interaction, and DNA binding [57].

Here, a total of 12 conserved motifs in the *G. arboreum*, *G. raimondii*, *G. barbadense* and *G. hirsutum* for *ERF* genes were identified. *Gene function diversity could be affected by* different numbers and types of motifs present in *ERF* proteins, in all the four species. Motif 1 and motif 2 in all *ERF* genes of cotton were highly conserved. The *ERF* family members in general shared similar structures of gene and motif compositions within same group, which proposes for the possibility of their similar roles in plant growth and development.

The study of expression patterns of genes is used to predicted the function of genes [49]. In this study the *ERF* genes showed high expression in ovule (0DPA, 1DPA, 3DPA, 5DPA, 10DPA, 20DPA) organs (root, stem, leaf apicalyx, receptacles), flower (sepal, petal, filament, anther) and fiber (10 DPA, 20 DPA, 25 DPA) across the species, suggesting that these genes have key roles in growth and development of cotton plant. Moreover, the expression of cotton *ERF* in different tissues or diverse stages, showed that these genes could be more stable than those that only expressed in specific tissues or one stage of an organ. Most of the *ERF* genes in *G. hirsutum* were also expressed in salt and heat stress. According to previous studies, some *ERF* genes were involved in various stresses responses in plants, such as high-salt, low-temperature and drought stress [42, 53]. Overall, the above findings provide foundation to further investigate the potential function of cotton *ERF* genes. These analyses are not only helpful in selecting valuable candidate *ERF* genes for further functional studies but also has important implication for genetic improvement for agricultural production and stress tolerance in cotton crop.

## 5. Conclusions

In this study, a genome-wide analysis of cotton *ERF* genes was performed. The 671 *ERF* genes in four species were identified and classified in detail. Protein lengths, molecular weights, and theoretical isoelectric points of cotton *ERF* vary greatly. The evolutionary characteristics, expression patterns of *ERF* genes in various cotton organs and growth stages, and their response to abiotic stress were studied. These results will be helpful to understand biological role of the *ERF* genes in cotton growth and development.

## Supplementary Materials

Supplementary materials can be found at www.mdpi.com/xxx/s1.

## Author Contributions

“Conceptualization, M.M.Z, A.R and A.R; methodology, A.R.; software, A.R; validation, A.R, AR and MMZ.; formal analysis, A.P.; investigation, G.M.; data curation, H.M.; writing—original draft preparation, MMZ; writing—review and editing, A.R and A.R; visualization, A.S, Y.Y, and M.R; supervision, and M.R; funding acquisition. All authors have read and agreed to the published version of the manuscript.”

## Ethics approval and consent to participate

Not applicable.

## Consent for publication

Not applicable.

## References

1. Loudet, O. and P.M. Hasegawa, Abiotic stress, stress combinations and crop improvement potential. The Plant Journal, 2017. 90(5): p. 837–838.

2. Cui, L., et al., Genome-wide identification, phylogeny and expression analysis of AP2/ERF transcription factors family in Brachypodium distachyon. BMC genomics, 2016. 17(1): p. 636.

3. Kaufmann, K. and C.A. Airoldi, Master regulatory transcription factors in plant development: a blooming perspective, in Plant Transcription Factors. 2018, Springer. p. 3–22.

4. Wessler, S.R., Homing into the origin of the AP2 DNA binding domain. Trends in plant science, 2005. 10(2): p. 54–56.

5. Xu, Z.S., et al., Functions and application of the AP2/ERF transcription factor family in crop improvement F. Journal of integrative plant biology, 2011. 53(7): p. 570–585.

6. Phukan, U.J., et al., Regulation of Apetala2/Ethylene response factors in plants. Frontiers in plant science, 2017. 8: p. 150.

7. Mantiri, F.R., et al., The transcription factor MtSERF1 of the ERF subfamily identified by transcriptional profiling is required for somatic embryogenesis induced by auxin plus cytokinin in Medicago truncatula. Plant Physiology, 2008. 146(4): p. 1622–1636.

8. Cai, X.-T., et al., Arabidopsis ERF109 mediates cross-talk between jasmonic acid and auxin biosynthesis during lateral root formation. Nature Communications, 2014. 5(1): p. 1–13.

9. Gu, C., et al., Multiple regulatory roles of AP2/ERF transcription factor in angiosperm. Botanical studies, 2017. 58(1): p. 1–8.

10. Ghorbani, R., et al., Genome-wide analysis of AP2/ERF transcription factors family in Brassica napus. Physiology and Molecular Biology of Plants, 2020. 26(7): p. 1463–1476.

11. Dossa, K., et al., Insight into the AP2/ERF transcription factor superfamily in sesame and expression profiling of DREB subfamily under drought stress. BMC plant biology, 2016. 16(1): p. 171.

12. Srivastava, R. and R. Kumar, The expanding roles of APETALA2/Ethylene Responsive Factors and their potential applications in crop improvement. Briefings in functional genomics, 2019. 18(4): p. 240–254.

13. Han, Z., et al., A genome-wide analysis of pentatricopeptide repeat (PPR) protein-encoding genes in four Gossypium species with an emphasis on their expression in floral buds, ovules, and fibers in upland cotton. Molecular Genetics and Genomics, 2020. 295(1): p. 55–66.

14. Fan, W.-B., et al., Comparative chloroplast genomics of Dipsacales species: Insights into sequence variation, adaptive evolution, and phylogenetic relationships. Frontiers in plant science, 2018. 9: p. 689.

15. Jin, L.-G. and J.-Y. Liu, Molecular cloning, expression profile and promoter analysis of a novel ethylene responsive transcription factor gene GhERF4 from cotton (Gossypium hirstum). Plant Physiology and Biochemistry, 2008. 46(1): p. 46–53.

16. Li, F., et al., Genome sequence of the cultivated cotton Gossypium arboreum. Nature genetics, 2014. 46(6): p. 567–572.

17. Wang, K., et al., The draft genome of a diploid cotton Gossypium raimondii. Nature genetics, 2012. 44(10): p. 1098–1103.

18. Zhang, T., et al., Sequencing of allotetraploid cotton (Gossypium hirsutum L. acc. TM-1) provides a resource for fiber improvement. Nature biotechnology, 2015. 33(5): p. 531–537.

19. Liu, X., et al., Gossypium barbadense genome sequence provides insight into the evolution of extra-long staple fiber and specialized metabolites. Scientific reports, 2015. 5: p. 14139.

20. Lamesch, P., et al., The Arabidopsis Information Resource (TAIR): improved gene annotation and new tools. Nucleic acids research, 2012. 40(D1): p. D1202–D1210.

21. Zhu, T., et al., CottonFGD: an integrated functional genomics database for cotton. BMC plant biology, 2017. 17(1): p. 1–9.

22. Singh, A.K., et al.,In silico analysis of FLP 18 gene of Anguina tritici and phylogenetic analysis with other nematodes.

23. Finn, R.D., et al., The Pfam protein families database: towards a more sustainable future. Nucleic acids research, 2016. 44(D1): p. D279–D285.

24. Finn, R.D., et al., HMMER web server: 2015 update. Nucleic acids research, 2015. 43(W1): p. W30–W38.

25. Yu, C.S., et al., Prediction of protein subcellular localization. Proteins: Structure, Function, and Bioinformatics, 2006. 64(3): p. 643–651.

26. Savojardo, C., et al., BUSCA: an integrative web server to predict subcellular localization of proteins. Nucleic Acids Research, 2018. 46(W1): p. W459–W466.

27. Sievers, F., et al., Fast, scalable generation of high_Jquality protein multiple sequence alignments using Clustal Omega. Molecular systems biology, 2011. 7(1): p. 539.

28. Kumar, S., G. Stecher, and K. Tamura, MEGA7: molecular evolutionary genetics analysis version 7.0 for bigger datasets. Molecular biology and evolution, 2016. 33(7): p. 1870–1874.

29. Chen, C., et al., TBtools: An Integrative Toolkit Developed for Interactive Analyses of Big Biological Data. Molecular Plant, 2020. 13(8): p. 1194–1202.

30. Wang, Y., et al., MCScanX: a toolkit for detection and evolutionary analysis of gene synteny and collinearity. Nucleic acids research, 2012. 40(7): p. e49–e49.

31. Dong, C., H. Hu, and J. Xie, Genome-wide analysis of the DNA-binding with one zinc finger (Dof) transcription factor family in bananas. Genome, 2016. 59(12): p. 1085–1100.

32. Xu, F.-C., et al., Heterogeneous expression of the cotton R2R3-MYB transcription factor GbMYB60 increases salt sensitivity in transgenic Arabidopsis. Plant Cell, Tissue and Organ Culture (PCTOC), 2018. 133(1): p. 15–25.

33. Jiang, Y., et al., Medicago AP2-domain transcription factor WRI5a is a master regulator of lipid biosynthesis and transfer during mycorrhizal symbiosis. Molecular plant, 2018. 11(11): p. 1344–1359.

34. Zhang, P., et al., The R2R3-MYB transcription factor MYB49 regulates cadmium accumulation. Plant physiology, 2019. 180(1): p. 529–542.

35. Fang, D.D., Cotton fiber: physics, chemistry and biology. 2018: Springer.

36. Müller, M. and S. Munné-Bosch, Ethylene response factors: a key regulatory hub in hormone and stress signaling. Plant physiology, 2015. 169(1): p. 32–41.

37. Li, X.-P., et al., Soybean DRE-binding transcription factors that are responsive to abiotic stresses. Theoretical and Applied Genetics, 2005. 110(8): p. 1355–1362.

38. Sharma, M.K., et al., Identification, phylogeny, and transcript profiling of ERF family genes during development and abiotic stress treatments in tomato. Molecular Genetics and Genomics, 2010. 284(6): p. 455–475.

39. Sharoni, A.M., et al., Gene structures, classification and expression models of the AP2/EREBP transcription factor family in rice. Plant and cell physiology, 2011. 52(2): p. 344–360.

40. Dietz, K.-J., M.O. Vogel, and A. Viehhauser, AP2/EREBP transcription factors are part of gene regulatory networks and integrate metabolic, hormonal and environmental signals in stress acclimation and retrograde signalling. Protoplasma, 2010. 245(1–4): p. 3–14.

41. Feng, J.-X., et al., An annotation update via cDNA sequence analysis and comprehensive profiling of developmental, hormonal or environmental responsiveness of the Arabidopsis AP2/EREBP transcription factor gene family. Plant molecular biology, 2005. 59(6): p. 853–868.

42. Sakuma, Y., et al., DNA-binding specificity of the ERF/AP2 domain of Arabidopsis DREBs, transcription factors involved in dehydration-and cold-inducible gene expression. Biochemical and biophysical research communications, 2002. 290(3): p. 998–1009.

43. Jiao, Y., et al., Ancestral polyploidy in seed plants and angiosperms. Nature, 2011. 473(7345): p. 97–100.

44. Li, F., et al., Genome sequence of cultivated Upland cotton (Gossypium hirsutum TM-1) provides insights into genome evolution. Nature biotechnology, 2015. 33(5): p. 524–530.

45. Cannon, S.B., et al., The roles of segmental and tandem gene duplication in the evolution of large gene families in Arabidopsis thaliana. BMC plant biology, 2004. 4(1): p. 10.

46. Zhu, Y., et al., Soybean (Glycine max) expansin gene superfamily origins: segmental and tandem duplication events followed by divergent selection among subfamilies. BMC plant biology, 2014. 14(1): p. 93.

47. Huang, Y., et al., Genome-wide identification and expression analysis of the ERF transcription factor family in pineapple (Ananas comosus (L.) Merr.). PeerJ, 2020. 8: p. e10014.

48. Li, P., et al., Genome-wide identification and expression analysis of AP2/ERF transcription factors in sugarcane (Saccharum spontaneum L.). BMC genomics, 2020. 21(1): p. 1–17.

49. Zhang, M., et al., Evolutionary and expression analyses of soybean basic Leucine zipper transcription factor family. BMC genomics, 2018. 19(1): p. 159.

50. He, D., et al., Genome-wide identification and analysis of the aldehyde dehydrogenase (ALDH) gene superfamily of Gossypium raimondii. Gene, 2014. 549(1): p. 123–133.

51. Lin, H., et al., Intron gain and loss in segmentally duplicated genes in rice. Genome biology, 2006. 7(5): p. R41.

52. Qiu, Y.-L., et al., The gain of three mitochondrial introns identifies liverworts as the earliest land plants. Nature, 1998. 394(6694): p. 671–674.

53. Nakano, T., et al., Genome-wide analysis of the ERF gene family in Arabidopsis and rice. Plant physiology, 2006. 140(2): p. 411–432.

54. Hu, L. and S. Liu, Genome-wide identification and phylogenetic analysis of the ERF gene family in cucumbers. Genetics and Molecular Biology, 2011. 34: p. 624–634.

55. Ahmed, S., et al.,Genome-wide investigation and expression analysis of APETALA-2 transcription factor subfamily reveals its evolution, expansion and regulatory role in abiotic stress responses in Indica Rice (Oryza sativa L. ssp. indica). Genomics, 2020.

56. Heyn, P., et al., Introns and gene expression: Cellular constraints, transcriptional regulation, and evolutionary consequences. BioEssays, 2015. 37(2): p. 148–154.

57. Liu, L., M.J. White, and T.H. MacRae, Transcription factors and their genes in higher plants. European Journal of Biochemistry, 1999. 262(2): p. 247–257.

